# POSITIVE RATE DEPENDENT ACTION POTENTIAL PROLONGATION BY MODULATING POTASSIUM ION CHANNELS

**DOI:** 10.1101/2022.03.06.483189

**Authors:** Candido Cabo

## Abstract

Pharmacological agents that prolong action potential duration (APD) to a larger extent at slow rates than at the fast excitation rates typical of ventricular tachycardia exhibit reverse rate dependence. Reverse rate dependence has been linked to the lack of efficacy of class III agents at preventing arrhythmias. In this report we show that, in computer models of the ventricular action potential, APD prolongation by accelerating phase 2 repolarization (by increasing IKs) and decelerating phase 3 repolarization (by blocking IKr and IK1) results in a robust positive rate dependence (i.e., larger APD prolongation at fast rates than at slow rates). In contrast, APD prolongation by blocking a specific potassium channel type results in reverse rate dependence or a marginal positive rate dependence. Interventions that result in a strong positive rate dependence tend to decrease the repolarization reserve because they require substantial IK1 block. However, limiting IK1 block to ∼50% results in a strong positive rate dependence with moderate decrease in repolarization reserve. In conclusion, the use of a combination of IKs activators and IKr and IK1 blockers could result in APD prolongation that potentially maximizes anti-arrhythmic effects (by maximizing APD prolongation at fast excitation rates) and minimizes pro-arrhythmic effects (by minimizing APD prolongation at slow excitation rates).

## INTRODUCTION

Many cardiac arrhythmias have a reentrant mechanism, a pattern of excitation in which a wave rotates around an anatomical or functional obstacle (Peters et al 2000). In order for the rotating wave to complete a reentrant cycle, the spatial extent of the reentrant wave (i.e., the wavelength, which is estimated as the product of myocardial conduction velocity times the tissue refractory period) has to be smaller than the perimeter of the anatomical or functional obstacle, leaving excitable tissue (i.e., an excitable gap) between the depolarizing head and the repolarizing tail of the reentrant wave. Class III pharmacological agents prolong the tissue refractory period by prolonging myocyte action potential duration (Vaughan-Williams 1975, Tamargo et al. 2004). The rationale behind the antiarrhythmic action of class III agents is that an increase in refractory period would lead to an increase in the wavelength of the reentrant wave with the consequent decrease in the excitable gap, causing the depolarizing head of the reentrant wave to run into refractory tissue extinguishing the arrhythmia.

However, clinical trials have shown that class III antiarrhythmic drugs are not effective in preventing initiation and maintenance of reentrant arrhythmias in post-myocardial infarction patients (Waldo et al. 1996, Bloch-Thomsen 1998, Køber et al. 2000). As it turns out, Class III agents prolong action potential duration to a greater extent at slow heart rates (for example during sinus rhythm) than at fast rates (for example during ventricular tachycardia), a property known as reverse rate dependence. This inability of class III agents to prolong action potential duration at the fast excitation rates typical of ventricular tachycardia has been linked to the lack of efficacy of class III agents at preventing arrhythmias (Hondeghem and Snyders 1990). Moreover, agents that prolong the action potential at slow rates may result in drug-induced LQT syndrome, trigger early afterdepolarizations and have a pro-arrhythmic effect (Kannankeril et al. 2010). Ideally, class III agents should prolong the action potential at fast rates with a minimal prolongation at slow rates, that is, they should exhibit a positive rate dependence, which is the opposite of a reverse rate dependence (Hondeghem and Snyders 1990; Winter and Shattock 2015).

It has been postulated that reverse rate dependence is an intrinsic property of ventricular myocardium (Banyasz et al. 2009). However, there is evidence that some pharmacological agents like amiodarone that prolong the action potential by blocking sodium and potassium channels have an attenuated reverse rate dependence response when compared to agents that block selectively a unique ion channel type (Hondeghem and Snyders 1990; Dorian and Newman 2000). We hypothesized that a prolongation of the action potential by modulating several potassium channels (instead of just a unique potassium channel type) may result in positive rate dependence or at least in an attenuated reverse rate dependence. We used computational models of the ventricular action potential in combination with optimization algorithms to investigate which interventions may result in prolongation of the action potential with a positive rate dependence (or an attenuated reverse rate dependence), and to further understand how the morphology of the action potential relates to its rate dependence.

## METHODS

### Computer models of the action potential

We simulated the cardiac action potential using the ten Tusscher-Noble-Noble-Panfilov (TNNP) computer model of a human ventricular epicardial cell (ten Tusscher et al. 2004) and the Hund-Rudy (HR) model of a canine ventricular cell (Hund and Rudy 2004). The models are publicly available and were downloaded from the CellML repository (www.cellml.org). We investigated the rate dependence of the action potential models by modulating the maximum conductance of the slow (IKs) and rapid (IKr) delayed rectifier potassium currents between 0 and 2x its standard value, and the maximum conductance of the inward rectifier potassium current (IK1) between 0.2 and 2x its standard value. All simulations were performed in a single cell. Action potentials were initiated with a depolarizing current with a strength 1.5x the stimulation threshold. We report measurements on action potentials that were calculated after 60 seconds of stimulation.

### Calculation of average total ionic current from the action potential shape

In a cell membrane, the total membrane current (*Im*) is the sum of the ionic current and the capacitive current:

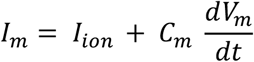

where *C*_*m*_ is the membrane specific capacitance (in μF/cm^2^) and *V*_*m*_ is the transmembrane voltage (in mV). For a single cell, there is no axial current, which means that *I*_*m*_ *= 0*, therefore:

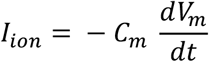

Integrating the previous equation from any two points in the action potential, V_m,t1_ (the membrane potential at t_1_) and V_m,t2_ (the membrane potential at t_2_), where t_2_ > t_1_, we obtain:

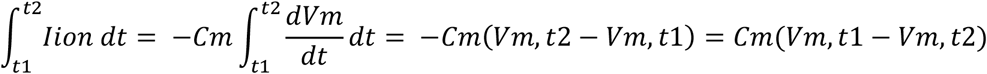

Using the mean value theorem, we can write:

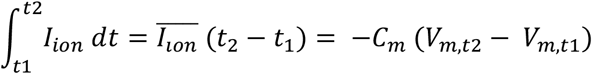

where 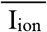 is the average of I_ion_ in the interval [t_1_, t_2_]. Ionic currents measured experimentally are typically normalized to the cell capacitance. Therefore, normalizing the previous expression to *C*_*m*_, we obtain:

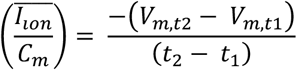

where now the current is in (pA/pF). This expression shows that the average 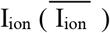 between any two points in the action potential is the slope of the line joining those two points.

### Action potential features

To quantify and compare different action potential shapes we extracted the following features from the action potential (Figure 1, top): action potential amplitude (APA); action potential duration at 90% repolarization (APD_90_); duration of phase 1, phase 2 and phase 3; average ionic current during phase 2 (I_ion, phase2_) and during phase 3 (I_ion, phase3_); ratio of I_ion, phase3_ and I_ion, phase2_; and the area under the action potential (AUAP) between the time of depolarization and the time of repolarization to APD_90_ normalized to APD_90_xAPA.

**Figure 1.**
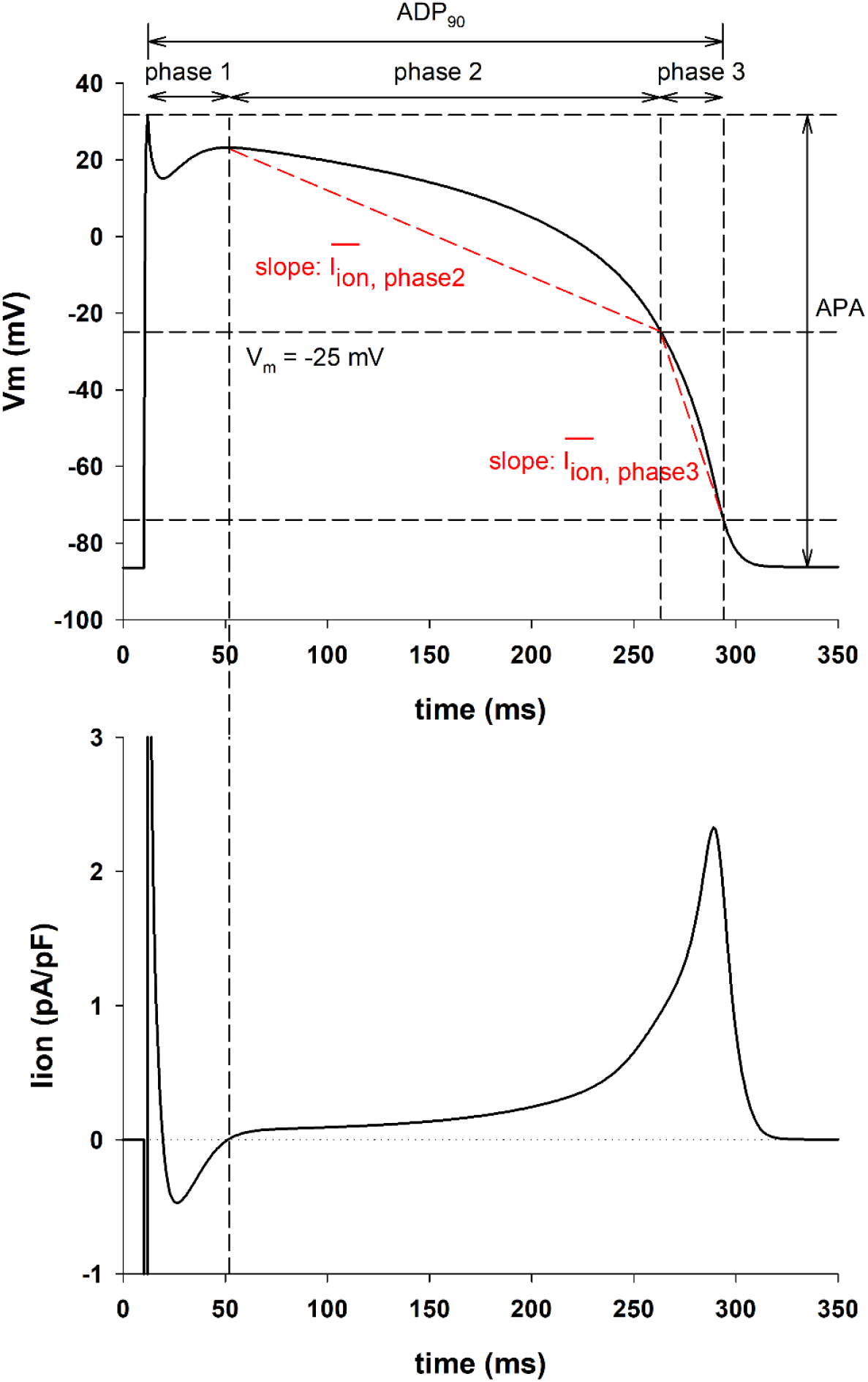
Features of the action potential (TNNP model). Top: Action potential calculated with the TNNP model with a BCL = 3000 ms. The figure shows the features of the action potential used in this study: action potential amplitude (APA); action potential duration at 90% repolarization (APD_90_); duration of phase 1, phase 2 and phase 3; average ionic current during phase 2 (I_ion, phase2_) and during phase 3 (I_ion, phase3_). Bottom: Total ionic current during the repolarization of the action potential. The vertical dashed line shows the time when Iion > 0 when phase 1 ends and phase 2 repolarization begins.

Phase 1 begins at the time of action potential depolarization and ends at the time repolarization starts, which is when the total ionic current (Figure 1, bottom) becomes positive (dashed vertical line in Figure 1). Phase 2 starts when phase 1 ends, and it ends when IK1 raises to 10% of its peak (we could not find in the literature a clear quantitative marker of the end of phase 2). In the TNNP model the end of phase 2 occurs when the membrane repolarizes to -25 mV. In the HR model the end of phase 2 occurs when the membrane repolarizes to -28 mV. Phase 3 starts at the end of phase 2, and it ends when the action potential repolarizes from its maximum depolarization potential to 90% of the action potential amplitude (APA).

The average ionic current during phase 2 (I_ion, phase2_ in Figure 1) is the slope of the line joining the V_m_ of the action potential where phase 2 starts and the point of the action potential where phase 2 ends (dashed red line in Figure 1). The same applies to the average ionic current during phase 3 (I_ion, phase3_ in Figure 1).

### Estimation of the repolarization reserve

Interventions that prolong APD may increase the risk of arrhythmias as a result of early afterdepolarizations. The concept of a repolarization reserve was developed to assess that pro-arrhythmic risk (Roden 1998). In this report, we estimated the repolarization reserve of a baseline action potential by quantifying the prolongation of the APD upon application of a constant depolarizing current of -0.1pA/pF during the action potential (Varro and Baczko 2011). This can be done experimentally for example by increasing the persistent sodium current with veratrine and anemonia sulcata toxin (ATX II) (Varro and Baczko 2011). With that protocol, a larger prolongation of the APD with respect to the baseline APD implies a smaller repolarization reserve and a higher risk of triggered arrhythmias.

### Optimization algorithm

When using potassium channel blockers to prolong the action potential, the prolongation of the action potential (with respect to control) at long cycle lengths (ΔAPD_long_) is generally larger than the prolongation of the action potential at short cycle lengths (ΔAPD_short_), resulting in reverse use dependence (ΔAPD_long_ > ΔAPD_short_). We used the Particle Swarm Optimization (PSO) algorithm (Kennedy and Eberhart, 1995) to find the optimal combination of conductance of IKs, IKr and IK1 to minimize the difference between ΔAPD_long_ and ΔAPD_short_. We used an implementation of the PSO algorithm publicly available in the Github repository (https://github.com/kkentzo/pso). Minimizing ΔAPD_long_-ΔAPD_short_ should result in an attenuation of reverse used dependence or achieving positive rate dependence if ΔAPD_long_ < ΔAPD_short_ for a given action potential duration. In the simulations presented here, the long cycle length was BCL = 3000 ms for both the TNNP and HR model, and the short cycle length was BCL = 400 ms for the TNNP model and BCL = 300 ms for the HR model. The PSO optimization algorithm allows to constraint the range of variation of the maximum conductance of the different potassium channels. In all PSO optimizer calculations, the maximum conductance of IKs and IKr was allowed to vary between 0 and 2x the control value. We run PSO optimizer calculations for different values of the lower limit of the maximum IK1 conductance (0.2x, 0.5x, 0.7x and 1x the control value) with an upper limit of 2x the control value.

## RESULTS

### Prolongation of the action potential with reverse and positive rate dependence

Figure 2A (top) shows the APD rate dependence during control (gray circles), and for four interventions that prolonged the control APD to 310 ms at BCL = 3000 ms, using the TNNP model. Note that while APD prolongation at BCL = 3000 ms (right vertical dashed line) is the same for all interventions, APD prolongation for shorter BCLs was markedly different. Figure 2A (bottom) shows the percentage APD prolongation with respect to control for the four interventions at different BCLs to illustrate better the reverse or positive rate dependence of the different interventions. APD prolongation by blocking IKs by 43% (0.57IKs, unfilled triangles down) shows reverse rate dependence because APD prolongation decreases with a decreasing BCL (from 9.5% at BCL = 3000 ms to 7% at BCL = 400 ms, vertical dashed lines). APD prolongation by 69% block of IKr (0.31IKr, unfilled triangles up) or 80% block of IK1 (0.2IK1, unfilled squares) shows a modest positive rate dependence (moderate increase in APD prolongation with decreasing BCL). On the other hand, the optimal combination of potassium channel activators (of IKs) and blockers (of IKr and IK1) found by the PSO algorithm (2IKs+0.33IKr+0.2IK1, black circles) shows a robust positive rate dependent response because APD prolongation increases with a decreasing BCL (from 9.5% at BCL = 3000 ms to 19% at BCL = 400 ms).

**Figure 2.**
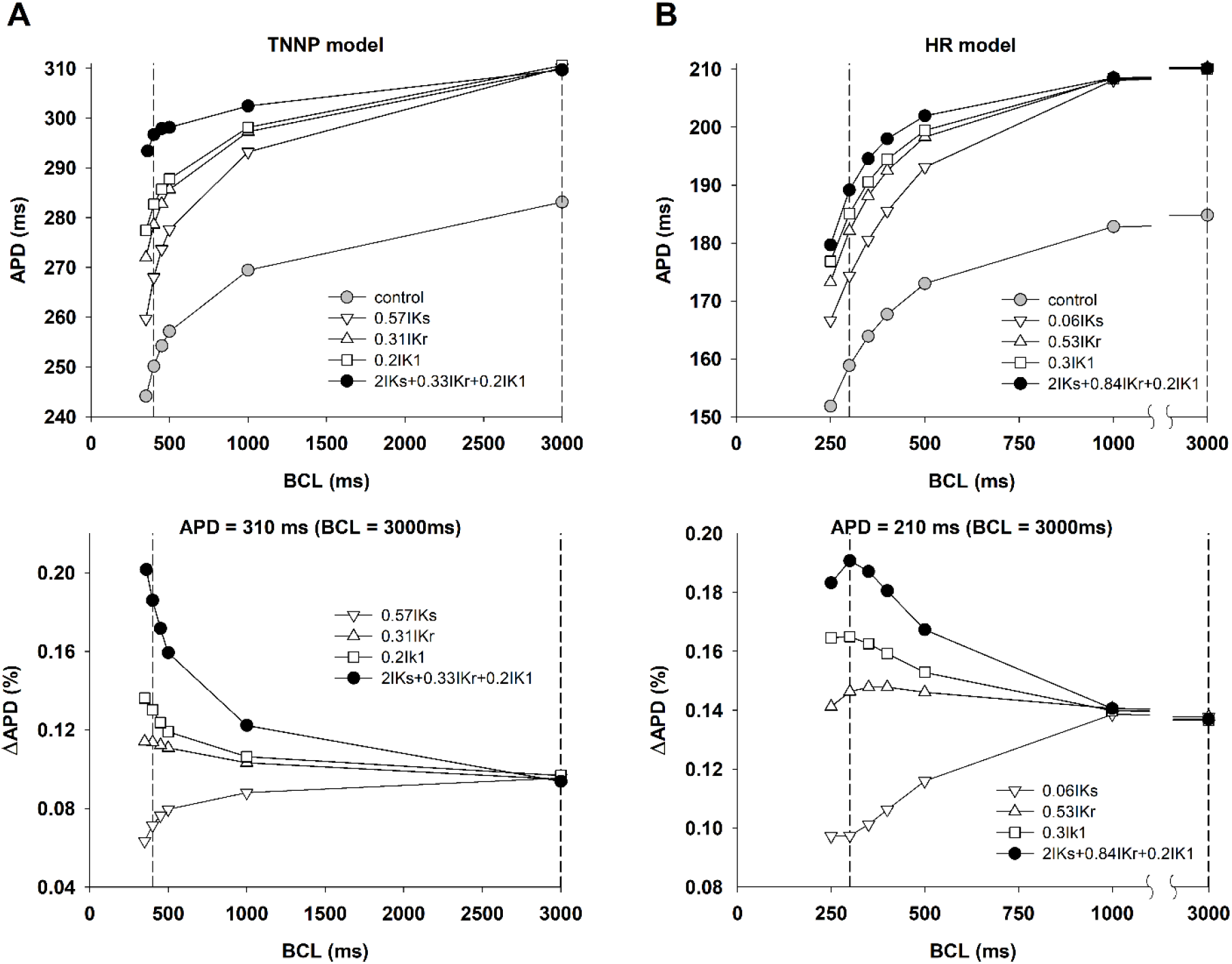
Panel A (top): APD rate dependence during control (gray circles), and for four interventions that prolonged the control APD to 310 ms when BCL = 3000 ms, using the TNNP model. The vertical dashed lines indicate BCL = 400 and 3000 ms respectively. Panel A (bottom): Percentage APD prolongation with respect to control for the four interventions in Panel A (top) at different BCLs to illustrate better the reverse or positive rate dependence of the different interventions. Panel B (top): APD rate dependence during control (gray circles), and for four interventions that prolonged the control APD to 210 ms at BCL = 3000 ms, using the HR model. The vertical dashed lines indicate BCL = 300 and 3000 ms respectively. Panel B (bottom): Percentage APD prolongation with respect to control for the four interventions in Panel B (top) at different BCLs to illustrate better the reverse or positive rate dependence of the different interventions. See text for detailed description.

Figure 2B shows similar results for interventions that prolong APD to 210 ms using the HR model. As with the TNNP model, while the prolongation of APD at BCL = 3000 ms (right vertical dashed line) is the same for all interventions, APD prolongation for shorter BCLs was markedly different (Figure 2B, top). As with the TNNP model, APD prolongation by blocking IKs (0.06IKs, unfilled triangles down) resulted in reverse rate dependence (Figure 2B, bottom), and APD prolongation with IKr block (0.53IKr, unfilled triangles up) or IK1 block (0.3IK1, unfilled squares) resulted in a moderate positive rate dependence. The optimal combination of potassium channel activators (of IKs) and blockers (of IKr and IK1) identified by the PSO algorithm (2IKs+0.84IKr+0.2IK1, black circles) again produced a stronger positive rate dependence than the interventions blocking a unique potassium channel type.

Figure 3A (top) summarizes the percentage APD shortening between BCL = 3000 ms and 400 ms (defined as (APD_BCL=3000_ – APD_BCL=400_) / APD_BCL=3000_) for different interventions that prolong the control APD to 295, 310 and 325 ms, using the TNNP model (vertical dashed-dot lines in Figure 3, top). Filled circles represent interventions that prolong ADP by modulating a unique potassium channel type; unfilled circles (labeled 1, 2, 3) represent interventions identified by the POS optimization algorithm to prolong APD maximizing positive rate dependence. Interventions along the central vertical dashed-dot line (APD = 310 ms) correspond to the interventions shown in Figure 2A. Note that the percentage APD shortening for interventions identified by POS optimization (unfilled circles) are always below 6% (horizontal dashed line), considerably smaller than for interventions that block a unique potassium channel type (filled circles).

**Figure 3.**
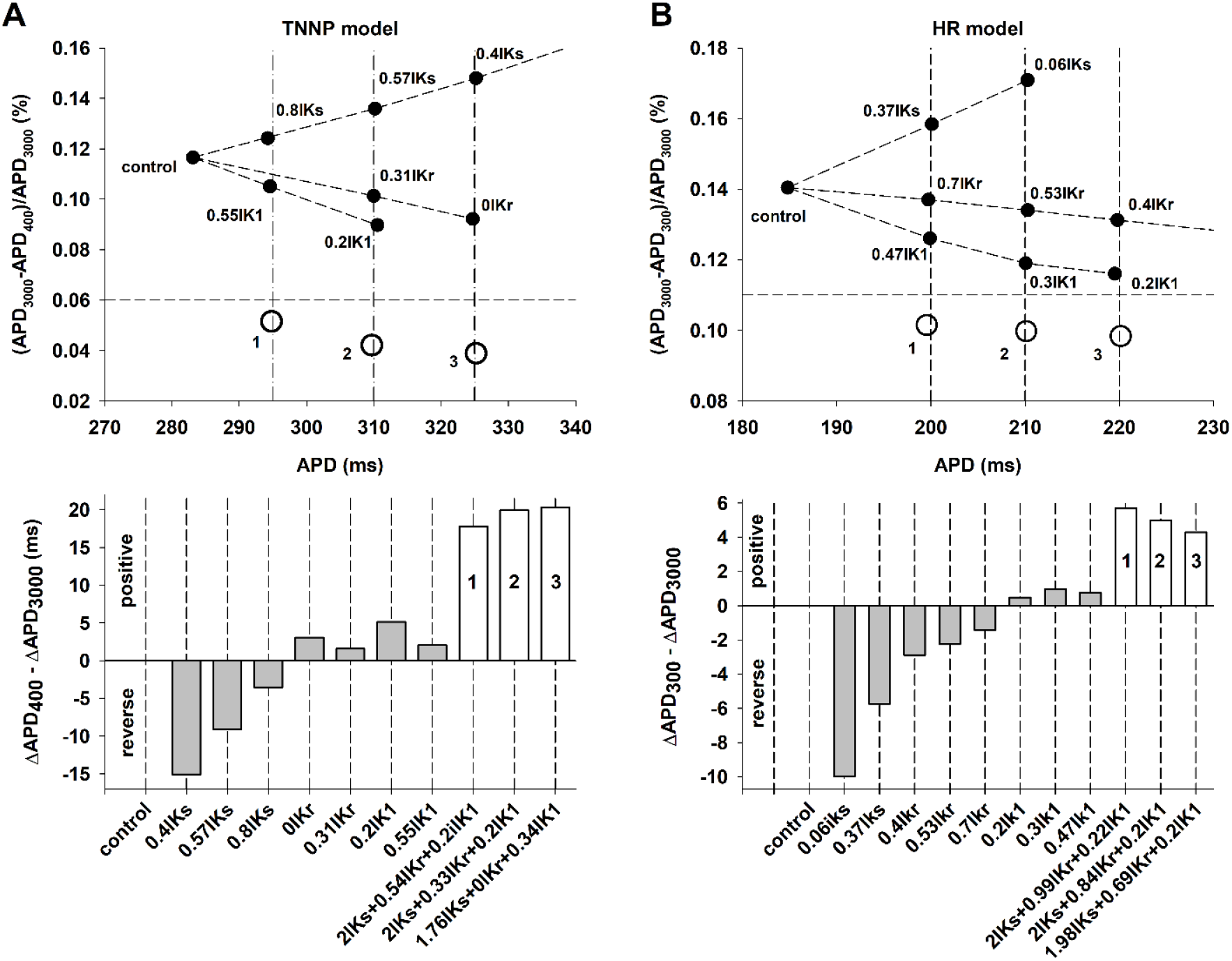
Panel A (top): Percentage APD shortening between BCL = 3000 ms and 400 ms, for different interventions prolonging APD by blocking a unique type of potassium channel (filled circles), and by activation/block of several potassium channels identified by POS optimization (unfilled circles), using the TNNP model. The vertical dashed-dot lines indicate different interventions that result in APDs of 295, 310 and 325 ms respectively when BCL = 3000 ms. Panel A (bottom): Difference between the prolongation of APD with respect to control for BCL = 400 ms and BCL = 3000 ms for the interventions in panel A (top). Negative values indicate reverse rate dependence and positive values indicates positive rate dependence. White bars labeled 1, 2, 3 indicate an intervention with a robust positive rate dependence and correspond to the unfilled circles in panel A (top). Gray bars indicate a reverse or marginal positive rate dependence. Panel B (top) Percentage APD shortening between BCL = 3000 ms and 300 ms for interventions that prolong APD to 200, 210 and 220 ms when BCL = 3000 ms (vertical dashed-dot lines), using the HR model. Filled/unfilled circles and labels have the same meaning as in Panel A (top). Panel B (bottom): Difference between the prolongation of APD with respect to control when BCL = 300 ms and BCL = 3000 ms for the interventions in panel B (top). Gray and white bars have the same meaning as in Panel A (bottom). See text for detailed description.

When APD prolongation (with respect to control) at BCL = 3000 ms (ΔAPD_BCL=3000_) is larger than APD prolongation (with respect to control) at BCL = 400 ms (ΔAPD_BCL=400_), i.e., when ΔAPD_BCL=400_ – ΔAPD_BCL=3000_ is negative, the intervention results in reverse rate dependence; otherwise, the intervention results in positive rate dependence. Figure 3A (bottom) shows whether the interventions in Figure 3A (top) exhibit a reverse (negative value in Figure 3A, bottom) or positive rate dependence (positive value in Figure 3A, bottom). Prolongation of APD by blocking independently IKs, IKr or IK1 results in either reverse rate dependence (for 0.4IKs, 0.57IKs and 0.8IKs) or a marginal positive rate dependence (0IKr, 0.31IKr, 0.2IK1 and 0.55IK1) (gray bars in Figure 3A, bottom). However, interventions that prolong the action potential by enhancing IKs and blocking IKr and IK1, result in a robust positive rate dependence response (white bars labeled 1, 2, and 3 in Figure 3A, bottom).

Figure 3B shows results of computations with the HR model, which are similar to those obtained with the TNNP model (Figure 3A). Interventions along the central vertical dashed-dot line (APD = 210 ms, Figure 3B, bottom) correspond to the interventions shown in Figure 2B. Again, note that the percentage APD shortening for interventions identified by POS optimization (unfilled circles) are always below 11% (horizontal dashed line), smaller than for interventions that block a unique potassium channel type (filled circles). Figure 3B (bottom) shows whether the interventions result in a reverse or positive rate dependence. As it happened with the TNNP model, interventions that prolong the action potential by enhancing IKs and blocking IKr and IK1, result in a robust positive rate dependence response (white bars labeled 1, 2, and 3 in Figure 3B, bottom).

Overall, the results in Figures 2 and 3 suggest that the TNNP model can produce a more robust positive rate dependence than that of the HR model. For example, ΔAPD_BCL=400_ – ΔAPD_BCL=3000_ is about 20 ms for the TNNP model (white bars, Figure 3A, bottom), but about 5 ms for the HR model (white bars, Figure 3B, bottom). However, for both models, for different amounts of APD prolongation, increasing IKs and blocking IKr and IK1 (interventions labeled 1, 2, 3 in Figures 3A and 3B) result in a robust positive rate dependence, which cannot be achieved by blocking a unique potassium channel type. Given that consistency, in what follows we present results of computations that use the TNNP model.

### Action potential shape influences its rate dependency

We then investigated how different features of the action potential shape influence its rate dependency. Figure 4A (top) shows the action potentials for three of the interventions (2IKs+0.54IKr+0.2IK1, 0.55IK1 and 0.8IKs) that resulted in APD = 295 ms with BCL = 3000 ms (vertical dashed-dot line through unfilled circle labeled 1 in Figure 3A, top), using the TNNP model. The action potentials are superimposed to make it easier to compare their shapes. Figure 4A (bottom) shows the corresponding total repolarizing currents during the action potential. Note that during the first 100 ms (phase 1 and beginning of phase 2) of the action potential there are no differences in shape, but the shape during phase 2 and phase 3 is markedly different for the three action potentials shown. The 2IKs+0.54IKr+0.2IK1 action potential (solid line in Figure 4A, top), repolarizes faster than the other two during phase 2, but slower during phase 3. This is also shown in Figure 4A (bottom) where the repolarizing current between 150 and 250 ms is largest for the solid line action potential, but during the final phase of repolarization (250 ms to 300 ms), the repolarizing current is largest for the 0.8IKs action potential (dashed-dot line in Figure 4A).

**Figure 4.**
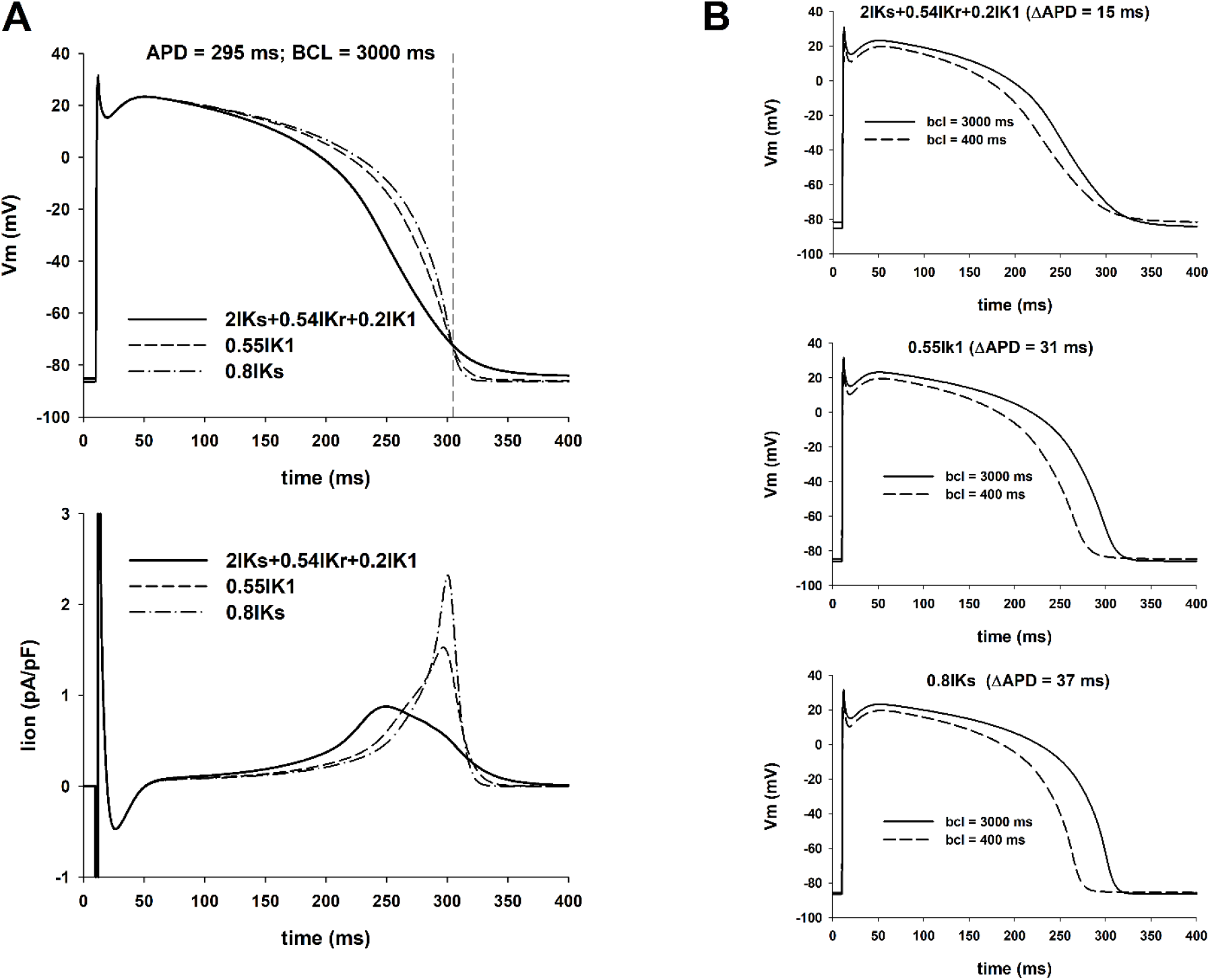
Influence of the action potential shape on its rate dependence (TNNP model). Panel A: Action potentials for different interventions that prolong the action potential to 295 ms for BCL = 3000 ms (top) with the corresponding total ionic currents (bottom) (vertical dashed-dot line labeled as 1 in Figure 3A, top). Panel B: Action potentials for BCL = 3000 (solid line) and 400 ms (dashed line) for the interventions in panel A. The action potential with a faster phase 2 repolarization and slower phase 3 repolarization (2IKs + 0.54IKr + 0.2IK1) results is a smaller APD shortening with increasing stimulation rate than the other two: 15 ms versus 31 ms for 0.55IKr and 37 ms for 0.8IKs.

Figure 4B shows rate dependent changes for the three action potentials in Figure 4A: solid action potentials were calculated with BCL = 3000 ms; dashed action potentials were calculated with BCL = 400 ms. The figure shows that the action potential that has a more pronounced phase 2 repolarization and less pronounced phase 3 repolarization (2IKs+0.54IKr+0.2IK1, Figure 4B, top) exhibits a smaller APD shortening (15 ms) with an increased stimulation rate than the other two action potentials (31 ms for 0.55IK1 and 37 ms for 0.8IKs).

Figures 5 and 6 shows similar results for APD = 310 ms and APD = 325 ms with BCL = 3000 ms (vertical dashed-dot lines through unfilled circles labeled 2 and 3 in Figure 3A, top). As in Figure 4, action potentials with a more pronounced repolarization during phase 2 and a less pronounced repolarization during phase 3 (resulting in a triangulation of the action potential) have a smaller APD percentage shortening with an increased stimulation rate, and result in a robust positive rate dependence (Figure 3A, bottom).

**Figure 5.**
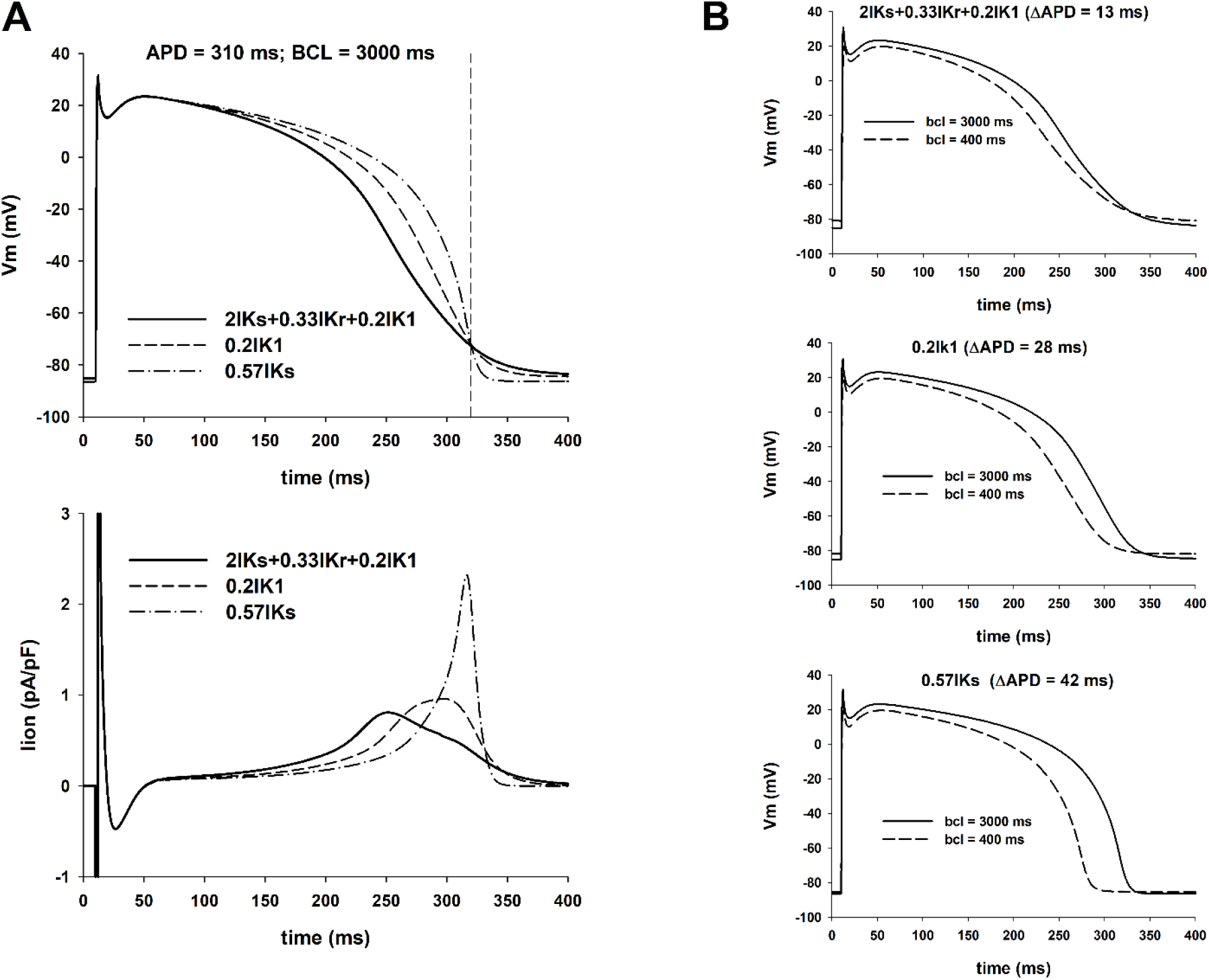
Influence of the action potential shape on its rate dependence (TNNP model). The figure shows action potentials and ionic currents resulting from interventions that prolong the action potential to 310 ms for BCL = 3000 ms (vertical dashed-dot line labeled as 2 in Figure 3A, top). The figure uses the same format as Figure 4.

**Figure 6.**
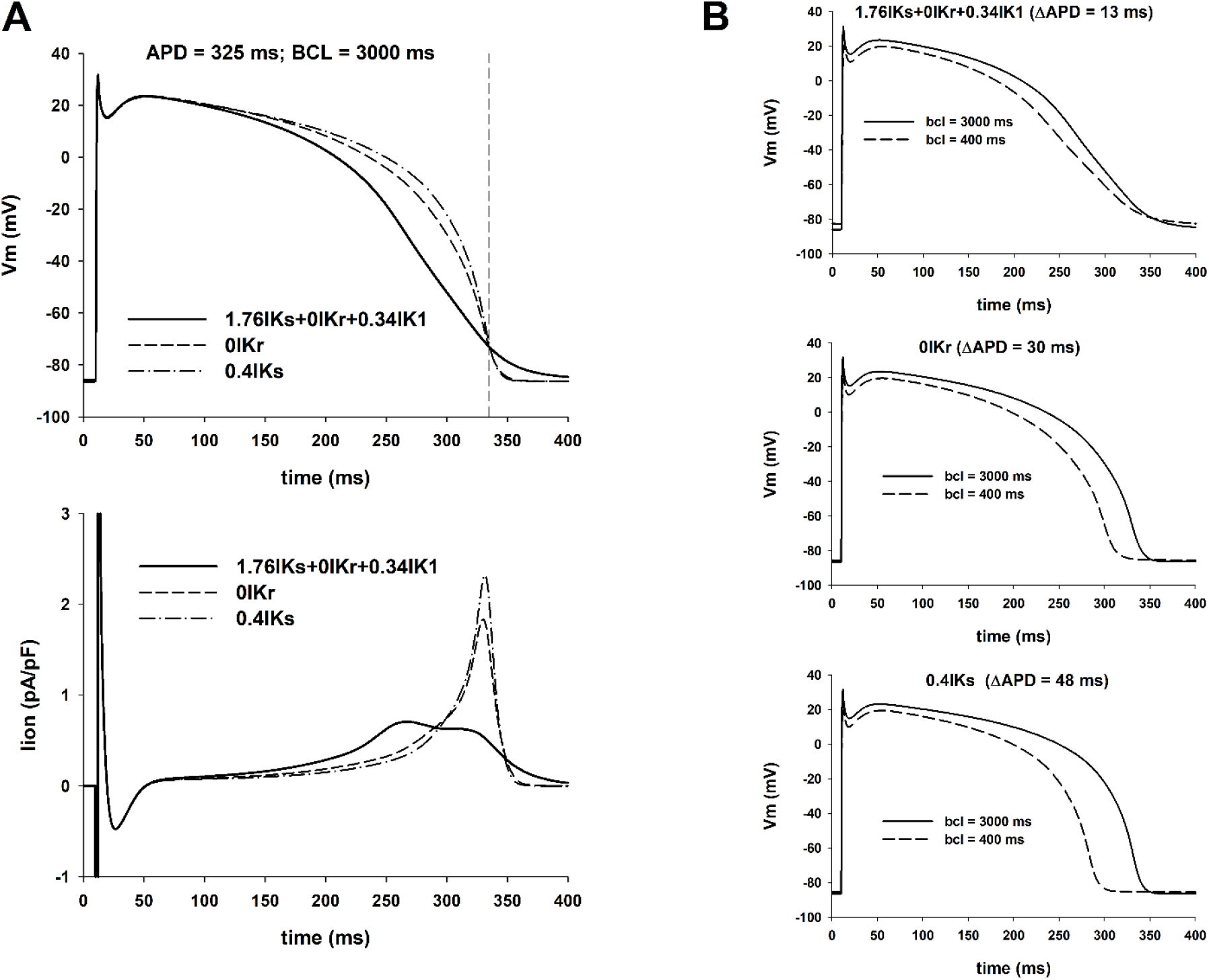
Influence of the action potential shape on its rate dependence (TNNP model). The figure shows action potentials and ionic currents resulting from interventions that prolong the action potential to 325 ms for BCL = 3000 ms (vertical dashed-dot line labeled as 3 in Figure 3A, top). The figure uses the same format as Figure 4.

### Action potential features and APD rate dependency

Figure 7 shows four features extracted from the action potentials in Figures 4, 5, and 6: average ionic current during phase 2, I_ion, phase2_ (panel A), average ionic during phase 3, I_ion, phase3_ (panel B), the ratio of the averages of I_ion, phase3_ and I_ion, phase2_ (panel C), and the area under the action potential (AUAP) between the time of depolarization and the time of repolarization to APD_90_ normalized to APD_90_xAPA (panel D). The white vertical bars represent features from action potentials found by the POS optimizer to prolong APD to 295, 310 and 325 ms with a maximal positive rate dependence response. Those action potentials result from the combined modulation of IKs, IKr and IK1. The gray vertical bars represent action potentials that prolong the APD to 310, 325 and 350 ms by blocking a unique potassium channel type and that result in reverse rate dependence (or a marginal positive rate dependence). Action potentials with a positive rate dependence are associated with a larger average ionic current during phase 2 (Figure 7A), a smaller average ionic current during phase 3 (Figure 7B), resulting in a smaller ratio of average ionic currents during phase 3 and phase 2 (triangularizing the action potential) and a smaller area under the action potential.

**Figure 7.**
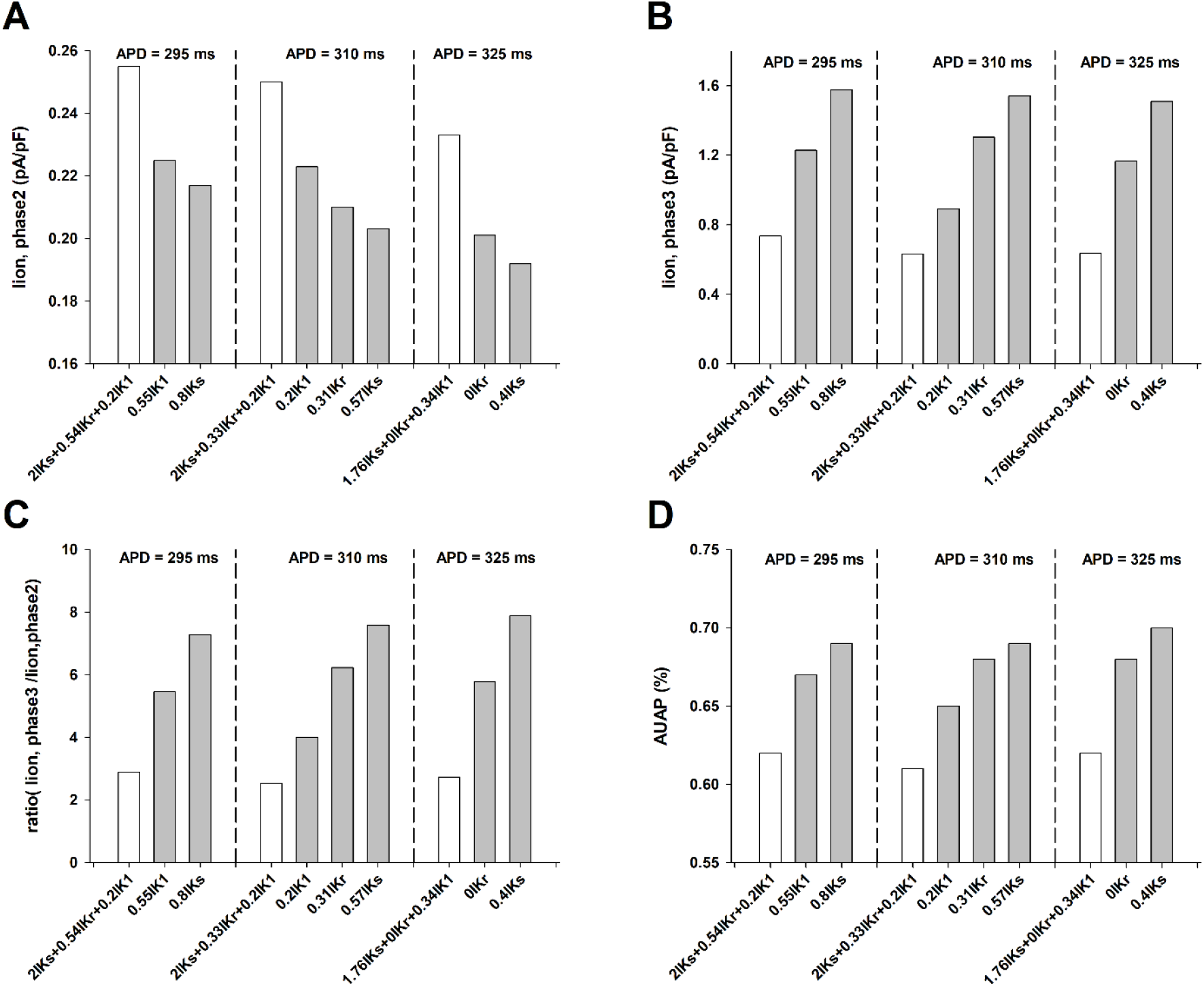
Features extracted from action potentials corresponding for the interventions in Figures 4 (APD = 295 ms), 5 (APD = 310 ms) and 6 (APD = 325 ms) for BCL = 3000 ms (TNNP model). White vertical bars represent interventions identified by the POS optimization algorithm, which involve the modulation of several potassium channels; gray vertical bars represent interventions involving modulation of one potassium channel. The features show the average ionic current during phase 2 (panel A), the average ionic current during phase 3 (panel B), the ratio of the average currents during phase 3 and phase 2 (panel C), and the area under the action potential (AUAP) normalized to APA and APD_90_ (panel D). The interventions identified by the POS algorithm result in a more triangularized action potential (smaller ratio in panel C and a smaller AUAP in panel D), with a positive rate dependence (Figure 3A, bottom).

The results in Figure 7 show that increasing phase 2 repolarization (for example by using IKs enhancers) and decreasing phase 3 repolarization (for example by using IKr and IK1 blockers) results in a smaller APD percentage shortening with an increased stimulation rate, and in a robust positive rate dependence.

### Effect of action potential triangulation on the repolarization reserve

Interventions that prolong the action potential with a positive rate dependence result in a triangulation of the action potential (Figure 7). It has been postulated that triangulation of the action potential could be pro-arrhythmic (Hondeghem et al 2001, Shah and Hondeghem 2005, Kannankeril et al 2010). In this section we estimate how the triangulation of the action potential affects the repolarization reserve (an indicator of the pro-arrhythmicity of an action potential) for the different interventions illustrated in Figures 4, 5, 6 and 7.

Figure 8 shows the percentage APD prolongation when a constant depolarizing current of - 0.1pA/pF is applied during the action potential for control (bar with the hatched pattern), and the different interventions that result in APDs of 295, 310 and 325 ms with BCL = 3000 ms. A larger APD prolongation with respect to the baseline APD indicates a smaller repolarization reserve. All interventions that prolong APD reduce the repolarization reserve when compared to control (horizontal dotted line in Figure 8). The interventions identified by the POS algorithm to prolong the APD with a positive rate dependence (white bars in Figure 8) decrease the repolarization reserve to a larger extent than interventions that prolong the APD with a reverse or marginal positive rate dependence (grey bars in Figure 8).

**Figure 8.**
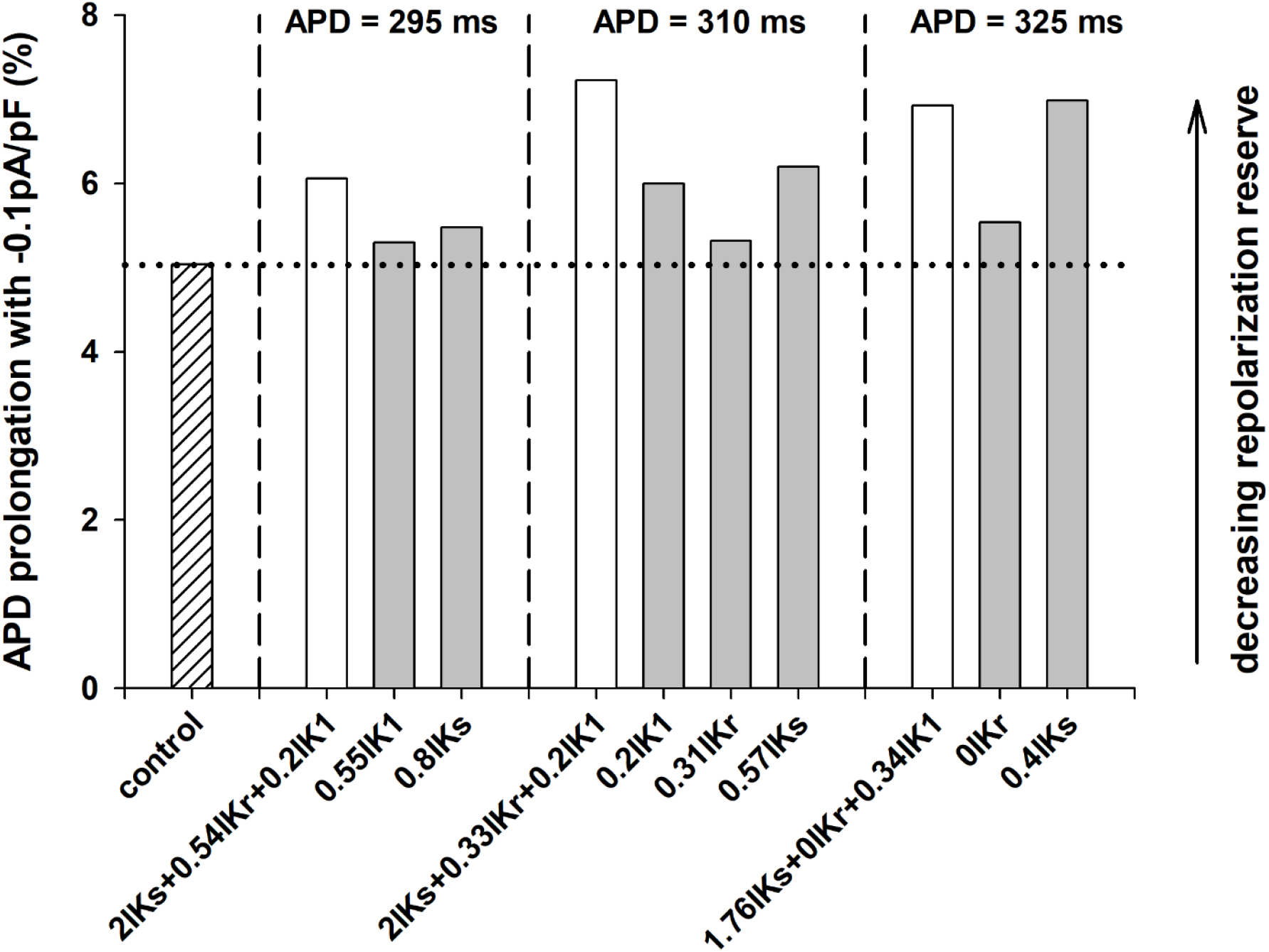
Percentage prolongation of the APD when a constant depolarizing current of -0.1pA/pF is applied during the action potential for control (bar with the hatched pattern), and the different interventions that result in APDs of 295, 310 and 325 ms at a BCL = 3000ms (TNNP model). A larger APD prolongation indicates a decreasing repolarization reserve. White bars indicate interventions with a robust positive rate dependence (see Figure 3A, bottom). Gray bars indicate interventions with reverse or marginal positive rate dependence (see Figure 3A, bottom).

### Effect of IK1 block on a positive rate dependence and the repolarization reserve

The substantial IK1 block (66%-80%) for the interventions that result in a positive rate dependent response (white bars in Figure 8) likely contributes to the decrease in the repolarization reserve. We then investigated if it is possible to achieve APD prolongation with a positive rate dependence with more moderate levels of IK1 block.

The PSO optimization algorithm allows to constraint the range of variation of the maximum conductance of the different potassium channels (see Methods). Figure 9 shows the results of the PSO optimization algorithm when the lower limit of the range of variation of the maximum IK1 conductance was 0.2x, 0.5x, 0.7x and 1x the value of control. In all cases the upper limit of the range was 2x the value of control. The range of variation of IKs and IKr remained between 0x and 2x the values of control.

**Figure 9.**
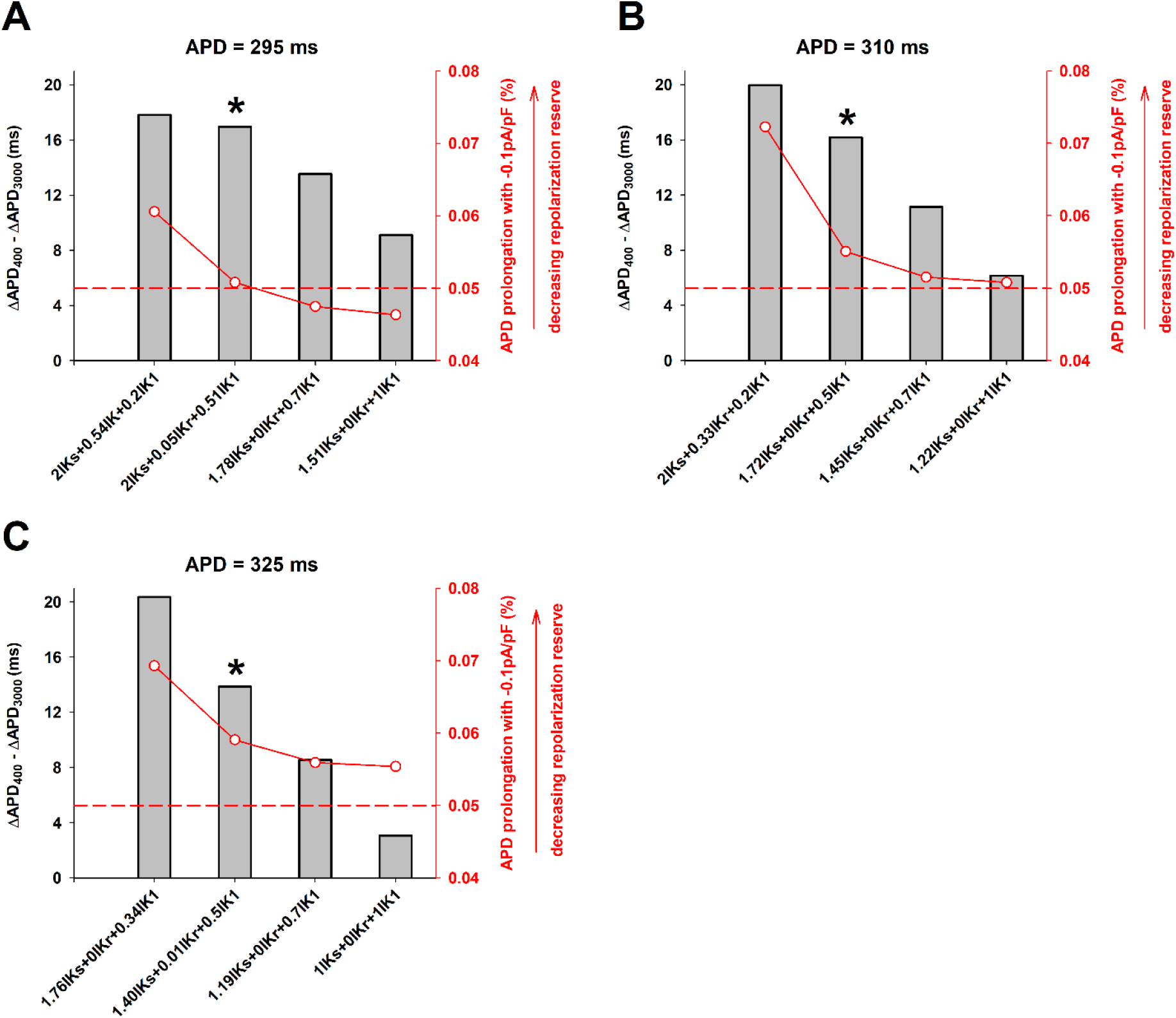
Interventions (x-axis) that result in a positive rate dependence response (gray bars, left y-axis) for different levels of IK1 block (80%, 50%, 30% and 0%) that result in APDs of 295 (panel A), 310 (panel B) and 325 ms (panel C) at a BCL = 3000ms (TNNP model). The right y-axis (red) shows the percentage prolongation of the APD when a constant depolarizing current of -0.1pA/pF is applied during the action potential. A larger APD prolongation indicates a decreasing repolarization reserve. The red horizontal dashed line shows the repolarization reserve for a control action potential. See text for explanation.

Figure 9A shows different interventions that result in a APD prolongation to 295 ms with a positive rate dependence (gray bars, left y-axis) for different levels of IK1 block (x-axis). Figure 9A also shows an estimation of the repolarization reserve for the different interventions (red plot, right y-axis). A higher value of the right y-axis indicates a smaller repolarization reserve and vice versa. The results indicate, as expected, that lower levels of IK1 block result in a larger repolarization reserve (monotonic decrease of the red line plot in Figure 9A). The repolarization reserve when IK1 block is between 0%-50% is the same or below that of a control action potential (horizontal red dashed line). Figure 9A also shows that more moderate levels of IK1 block result in a decreasing positive rate dependence response.

Figures 9B and 9C show similar results for interventions that prolong APD to 310 and 325 ms respectively. The strength of the positive rate dependence response and the repolarization reserve go in different directions: a stronger positive rate dependence response results in a decreasing repolarization reserve and vice versa. However, most of the decrease in the repolarization reserve occurs for interventions that block IK1 by more than 50%. Interventions that limit IK1 block to 50% still result in a robust positive rate dependence response with a repolarization reserve similar to that of a control action potential (grey bars with an asterisk in Figure 9A, 9B and 9C).

## DISCUSSION

We have shown that, in computer models of the ventricular action potential, APD prolongation by accelerating phase 2 repolarization (by increasing IKs) and decelerating phase 3 repolarization (by blocking IKr and IK1) results in a robust positive rate dependence. In contrast, APD prolongation by blocking a specific potassium channel type results in reverse rate dependence or a marginal positive rate dependence. Interventions that result in a strong positive rate dependence generally decrease the repolarization reserve because they require substantial IK1 block. Limiting IK1 block to ∼50% could still result in a strong positive rate dependence with moderate decrease in repolarization reserve.

There is abundant experimental evidence showing that APD prolongation by blocking a unique potassium channel type results in reverse rate dependence (Virag et al. 2009, Banyasz et al. 2009, Barandi et al 2010). Banyasz et al. (2009) and Barandi et al. (2010) showed consistent reverse rate dependence with several interventions causing lengthening and shortening of the action potential and concluded that reverse rate dependence is an intrinsic property of ventricular myocardium. Clinical results show that APD prolongation by blocking IKr in humans, with different channel specific pharmacological agents, also results in reverse rate dependence (Table 1 in Dorian and Newman 2000). Thus, the available experimental and clinical evidence suggest that it is not possible to prolong APD with a positive rate dependence response by blocking a specific potassium channel type. That is consistent with the results of the computations presented in this report (Figures 2 and 3).

On the other hand, there is clinical and experimental evidence suggesting that it is possible to achieve APD prolongation with a neutral (neither reverse nor positive) rate dependence by modulating several ion channels, either by using one agent that interacts with different ion channels or by using several agents each one interacting with a specific ion channel type. For example, amiodarone, an agent that prolongs APD by blocking sodium and potassium channels, has an attenuated reverse rate dependent response when compared to agents that block selectively a unique ion channel type (Hondeghem and Snyders 1990; Dorian and Newman 2000). A reduction in reverse rate dependence can also be achieved using a combination of drugs affecting several channel types. For example, in an experimental model of the canine infarcted heart, the combination of esmolol, a short acting beta-blocker that blocks IK1 and the L-type Ca channel (Shibata et al. 2012), and d-sotalol (an IKr blocker) prevented initiation of sustained ventricular tachycardia and attenuated the reverse rate dependence of d-sotalol (Kassotis et al. 2003). It is possible that the mechanism for the absence of reverse rate dependence during treatment with esmolol and d-sotalol is similar to the mechanism causing a positive rate dependence response in the computations presented here. Block of the L-type Ca channel by esmolol (Shibata et al. 2012) could have resulted in an acceleration of phase 2 repolarization, and IK1 block by esmolol (Shibata et al. 2012) and IKr block by d-sotalol (Tamargo et al. 2004) could have caused deceleration of phase 3 repolarization. Table 1 in Dorian and Newman (2000) summarizes the effects of class III antiarrhythmic drugs on rate dependence in humans: amiodarone and d-l-sotalol (combination of d-sotalol and a beta-blocker) show a neutral rate-dependence, while different specific IKr blockers are all reverse rate dependent. Overall, these results suggest that the use of a combination of agents that affect various channels could be effective in achieving at least an attenuated reverse rate dependence with a consequent antiarrhythmic effect (Hondeghem and Snyders 1990; Dorian and Newman 2000).

In that context, we investigated optimal combinations of potassium channel activators and blockers that could result in a robust APD prolongation with positive rate dependence, and evaluated their potential pro-arrhythmic risk. Our simulations, using two different computer models of the ventricular action potential (ten Tusscher et al. 2004, Hund and Rudy 2004), consistently show that interventions that increase IKs and block IKr and IK1 prolong APD with positive rate dependence (Figures 2 and 3). But those interventions, which triangularize the action potential (Figure 7), could potentially decrease the repolarization reserve and be pro-arrhythmic (Hondeghem et al 2001, Shah and Hondeghem 2005, Kannankeril et al 2010). Figure 9 shows that interventions with higher levels of IK1 block (resulting in a more extreme action potential triangulation) lead to stronger positive rate dependence responses at the cost of a larger decrease in the repolarization reserve (Figure 9). However, interventions with moderate IK1 block (<= 50%) still result in a robust positive rate response with moderate decrease in the repolarization reserve with respect to control (Figure 9, horizontal red dashed line).

All interventions that result in a positive rate dependence response in our computations, for different amounts of APD prolongation and for different levels of IK1 block, require the use of IKs activators (Figure 9, x-axis in panels A, B and C). IKs activators have been developed to prevent excessive APD prolongation that may occur in patients suffering LQT syndrome, cardiac hypertrophy or cardiac failure (Xu et al. 2002, Tamargo et al. 2004, Xu et al. 2015). IKs activators may also prevent excessive APD prolongation resulting from IKr and IK1 block in our simulations, and contribute to the modest decrease of the repolarization reserve for interventions with positive rate dependent response (Figure 9). All in all, it is possible that the use of a combination of IKs activators and IKr and IK1 blockers could result in APD prolongation that potentially maximizes anti-arrhythmic effects (by maximizing APD prolongation at fast rates) and minimizes pro-arrhythmic effects (by minimizing APD prolongation at slow rates) (Hondeghem and Snyders 1990).

Several hypotheses have been proposed to explain the mechanism of reverse rate dependent APD prolongation by blocking potassium channels (Barandi et al 2010). Incomplete deactivation of IKs at fast rates could result in a repolarizing force that counteracts the effect of IKr block to prolong APD at fast rates (Jurkiewicz and Sanguinetti, 1993). It is also possible that an increase in the extracellular K^+^ concentration at fast rates could hinder the APD prolongation efficacy of IKr block at fast rates (Yang and Roden 1996). Ion channels open and close during the cardiac cycle, and the interaction of ion channels and pharmacological agents may depend on the state (open or close) of the channel (Hondeghem and Katzung, 1977). During fast rates, myocardium is depolarized during most of the cardiac cycle (the opposite is true at slow rates). Therefore, if a pharmacological agent binds to the close state of the channel (i.e., during diastole) and unbinds when the channel is open (i.e., during systole), it will be less effective at fast rates than at slow rates, which would explain APD prolongation with reverse rate-dependence. The repolarization rate during late repolarization (i.e., the average I_ion_ current during phase 3 in this report, see Methods) can also affect the dynamics of IKr and IK1 (which in turn will also affect the repolarization rate) and have implications for reverse rate dependency (Virag et al. 2009). More recently, it has also been proposed that reverse rate dependence is an intrinsic property of ventricular myocardium resulting from the non-linear relationship between the rate of repolarization (I_ion_) and APD (Banyasz et al. 2009, Barandi et al. 2010, Winter and Shattock 2015). A consequence of that non-linear relationship would be that baseline APD may be a determining factor contributing to reverse rate dependence (Barandi et al 2010). Still, our results show that action potentials having the same APD but different shapes can have drastically different rate dependent responses (Figures 4, 5, 6). Therefore, it is likely that there are multiple factors contributing to reverse (and positive) rate dependence, and our results suggest that the shape of the action potential could be one of those factors (Figure 7).

### Limitations

A number of factors should be considered when interpreting the results presented in this report. Computer models of the action potential have inherent limitations because model parameters are usually estimated from experimental data obtained under different conditions and from different preparations. Moreover, sometimes the experimental data needed to formulate the models is contradictory and/or incomplete. Despite those limitations and the differences in ion channel density and kinetics between the TNNP and HR models, both models predict APD prolongation with a positive rate dependence for interventions that enhance IKs and block IKr and IK1 (Figures 2 and 3).

The density and kinetics of ion channels of myocytes is typically altered by disease. For example, myocardial infarction leads to the remodeling of several ion currents (Cabo and Boyden 2003), which increases the heterogeneity of ventricular myocardium and creates a substrate that makes it possible to initiate and sustain ventricular tachycardia (Baba et al. 2005). The effect of drug agents on remodeled myocytes may be different from their effect in healthy myocytes (Cabo and Boyden 2003). Therefore, whether or not enhancing IKs and blocking IKr and IK1 will result in APD prolongation with a positive rate dependence in the remodeled myocardium that provides the substrate for ventricular tachycardia is unknown and deserves further study.

In addition to ion channel remodeling, gap junction conductance and distribution is also remodeled by disease (Cabo et al. 2006). Therefore, even if APD prolongation with positive rate dependence in remodeled myocytes is possible, it is unknown whether that intervention will be effective in preventing initiation and/or maintenance of ventricular tachycardia in heterogeneous myocardium. Further computational and experimental studies will be needed to investigate how interventions that cause APD prolongation with positive rate dependence in single cells affect the dynamics of propagation of premature impulses that initiate and sustain reentrant arrythmias.

